# Effects of the secondhand smoking exposure in the early stages of the bone development

**DOI:** 10.1101/553834

**Authors:** Cristiano Fittipaldi Alves, Cesar Alexandre Fabrega Carvalho, Antônio Francisco Iemma, Francisco Haiter Neto, Paulo Henrique Ferreira Caria

## Abstract

**Objective:** The objective of this study was to evaluate the effects of the secondhand smoking in the trabecular bone micro-architecture of the mandible of rats, offsprings of passive smoking matrices.

**Materials and Methods:** Fifty-five rats, *Rattus norvegicus albinus*, offsprings of passive smoking and non-passive smoking matrices, were divided into three groups: continuous smoking offsprings (CSO), interrupted smoking offsprings (ISO) and non-smoking offsprings (NSO/control). After the 21^st^, 42^nd^, 63^rd^ and 128^th^ days, the mandibles were analyzed by micro-computer tomography(micro-CT). Images of inter-radicular alveolar bone of the mandibular first molars underwent three-dimensional reconstruction and were analyzed. The bone volume fraction (BV/TV, bone volume/total volume), the trabecular thickness (Tb.Th), the trabecular spacing (Tb.Sp), the trabecular number (Tb.N) and the structure model index (SMI) were analyzed.

**Results:** The BV/TV analysis revealed increase of the average values in the CSO group, at 21^st^ and 42^nd^ days (p=0,0124), tending to decrease related to the mean from the 42^nd^ day. The animals of ISO group did not show significant difference in BV/TV, about the control group (p=0,9751). The results of Tb.Th were different and significant during all the experimental period among the three groups: CSO and control (p<0,0001), ISO and control (p=0,0030) and CSO / ISO (p=0,0020). About Tb.Sp, the differences were not significant among the three groups. About Tb.N, the difference was significant into each group, with increasing values (p<0.0001). The SMI showed significant difference between the CSOs and control, CSO and ISO, both with (p<0,0001). The difference between control and ISO group was not significant (p=0,1253).

**Conclusion:** The passive inhalation of cigarette smoke by the offsprings of smoking matrices had a harmful effect in the micro-archicteture of the trabecular bone of the rats’ mandible in developing. About the ISO groups, the recovery of the micro-archicteture occurred partially.

## Introduction

There is an estimated that 40% of the children in the world, 33% of boys and 35% of girls, are exposed to the secondhand smoke. [1]. The cigarette smoke inhaled in a passive way consists of 15% of the mainstream smoke, the smoke inhaled by the current smoker and 85% of the sidestream smoke. The sidestream smoke, coming from the lighted cigarette, is considered more toxic than the mainstream smoke (inhaled)[2]. The secondhand smoke is considered the third largest cause of avoidable death worldwide and the main polluting agent in indoor environments [3]. The harmful effects to health due to the secondhand smoke were observed in current smokers’ wives in a study that lasted 14 years. [4]. The cigarette smoke toxicity reduces the blood nourishment and it can causes a cell division reduction, DNA damage induction, increased wound healing time and decreased tissue regenerative capacity.[5-6]. Such reduction causes permanent artery damages [7] that increase the risk of the disease development and alterations such as osteoporosis [8-9]. The osteoporosis is an osteometabolic disorder [10] characterized by the bone mass loss and by microstructure alterations which cause weakness and fracture risk [11-12]. The nicotine in cigarretes suppresses the function of the red blood cells, causes vasoconstriction and reduces the blood oxygen levels [13], and it reduces the calcium and estrogen levels as well [14], important substances in the bone growth and maintenance [15]. The nicotine also modulates several factors of the osteoblasts differentiation, proliferation and growth and it can decrease during the bone neoformation, osteogenic expression, [16-17], healing and fracture regeneration [18]. The effects caused by the cigarette smoke toxicity, inhaled in a passive way during the bone remodelling, suggest the reduction of the bone mineral density and the alteration in the microstructural bone organization in passive smokers. [19]. The concept of the bone quality evolved from an approach only based on the density to a structural micro-architecture [20]. In a recent study it was showed that the trabecular bone weakness and the bone mineral content indicate the increase of the fracture risk. [21]. However, there are controversies related to the possibility and the time of the bone tissue recovery before the adverse effects of the nicotine passive exposition [22-23]. Thus, the micro-computed tomography is an effective tool when evaluating the cigarette smoke influence in the bone sctructure[24]. It is an accurate, fast and non-destructive method, and it allows the measurement of the microstructures in non-processed biopsies and even in small bones, establishing automatically tridimensional histomorphometric rates [25].

Considered the reasons here presented, the purpose of this study was to evaluate the cigarette smoke toxic effects in the mandibular trabecular bone micro-architecture in developing, and its possible recovering after the cessation of the inhalation, in passive smoker rats which were offsprings of smoking matrices, in different periods.

## Materials and methods

### Treatment of the matrices

This study was approved by an Ethics Committee for the Animals Use of Faculty of Medicine of Jundiaí, number 285/2013). The matrices of the offsprings were composed of twenty *Rattus norveegicus albinus (Wistar)*, at the age of 8 weeks, and weighing 340 ± 10g. There were four male rats and four female rats in the control group, and there were six male rats and six female rats in the smoking group. There were two rats in each cage, kept at a room temperature (22+/-2°C) and in a light/dark-12 hours-cycle, with water and feed *ad libitum*.

### Exposure to the cigarette smoking and formation of the experimental groups

It was used an easy commercially avaliable cigarette due to the adaptation to smoke (tar: 10mg, and nicotine: 0,8mg). The matrices were gradually exposed to the cigarette smoke until to complete a total of 20 cigarettes units per day, in a period of eight weeks. Then, couples were formed with the mating purpose. The lighted cigarettes were put in the influx system of a vented shelf (fig 1) and, in an homogeneous way, the smoke of the lateral flux was spread into the sealed chambers [24]. During all the gestation, neonatal, breastfeeding, and growth period, the offsprings of passive smoking matrices and the matrices, were exposed to the smoke. After the weaning, 35 male offsprings were selected and kept under the smoke exposition at the same initial conditions of their mothers, constituting 4 groups of continuous smoking offsprings (CSO). The offsprings submitted to the cessation of the inhalation – the group of interrupted smoking offsprings (ISO), were formed from the CSO subdivision in a period equivalent to half of the exposure time to the cigarette smoke (Fig 2). It was not formed a 21day-ISO-group because it was a breastfeeding period, and the milk abstinence would be harmful to the bone development of the offsprings. The non-smoking offsprings (control group or NSO) were kept in a cigarette smoke-free exposure environment (Fig 2).

**Fig 1.**
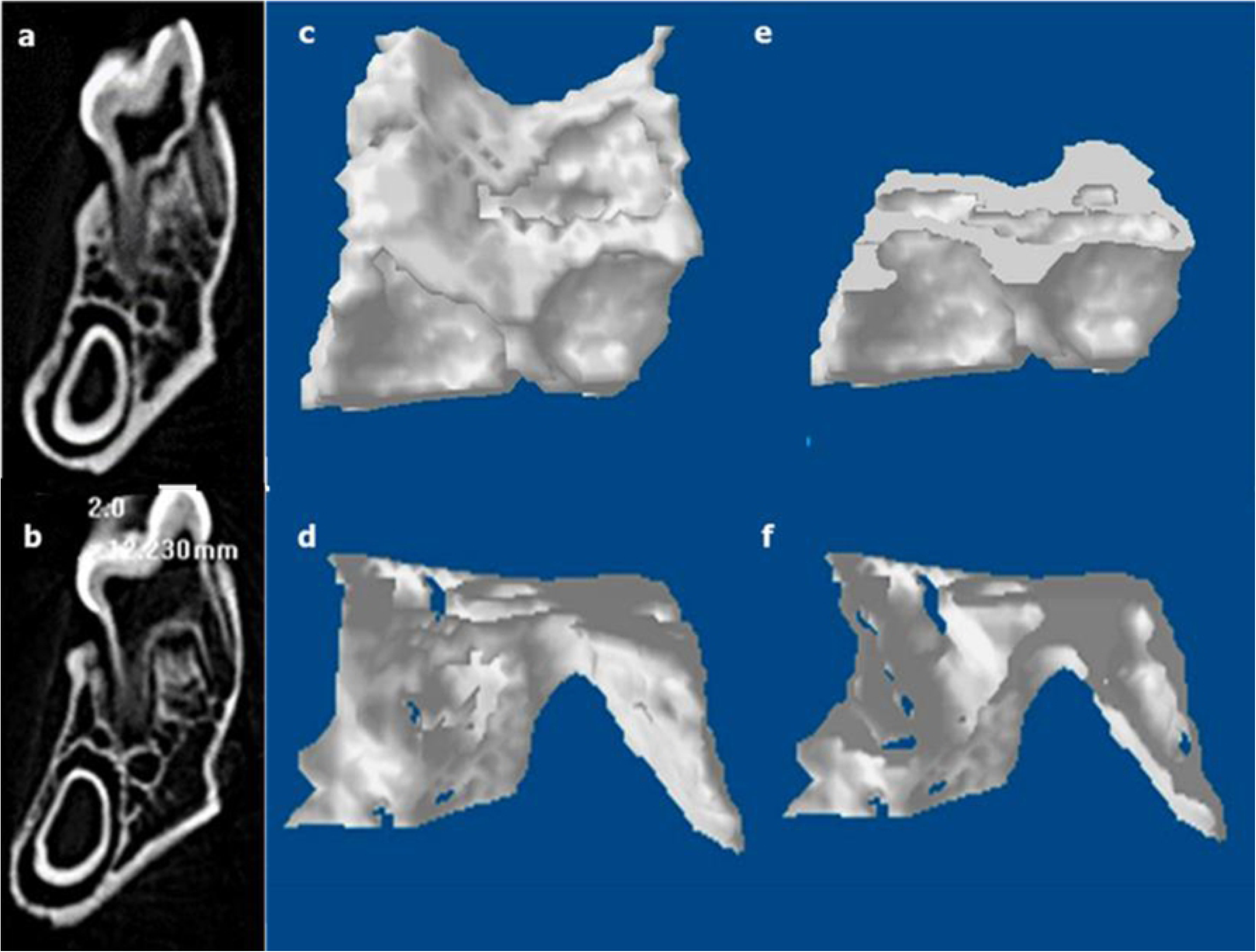
Vented shelf. a: frontal aspect, b: cigarettes put in the area of the influx of the smoke for the sealed chambers.

**Fig 2.**
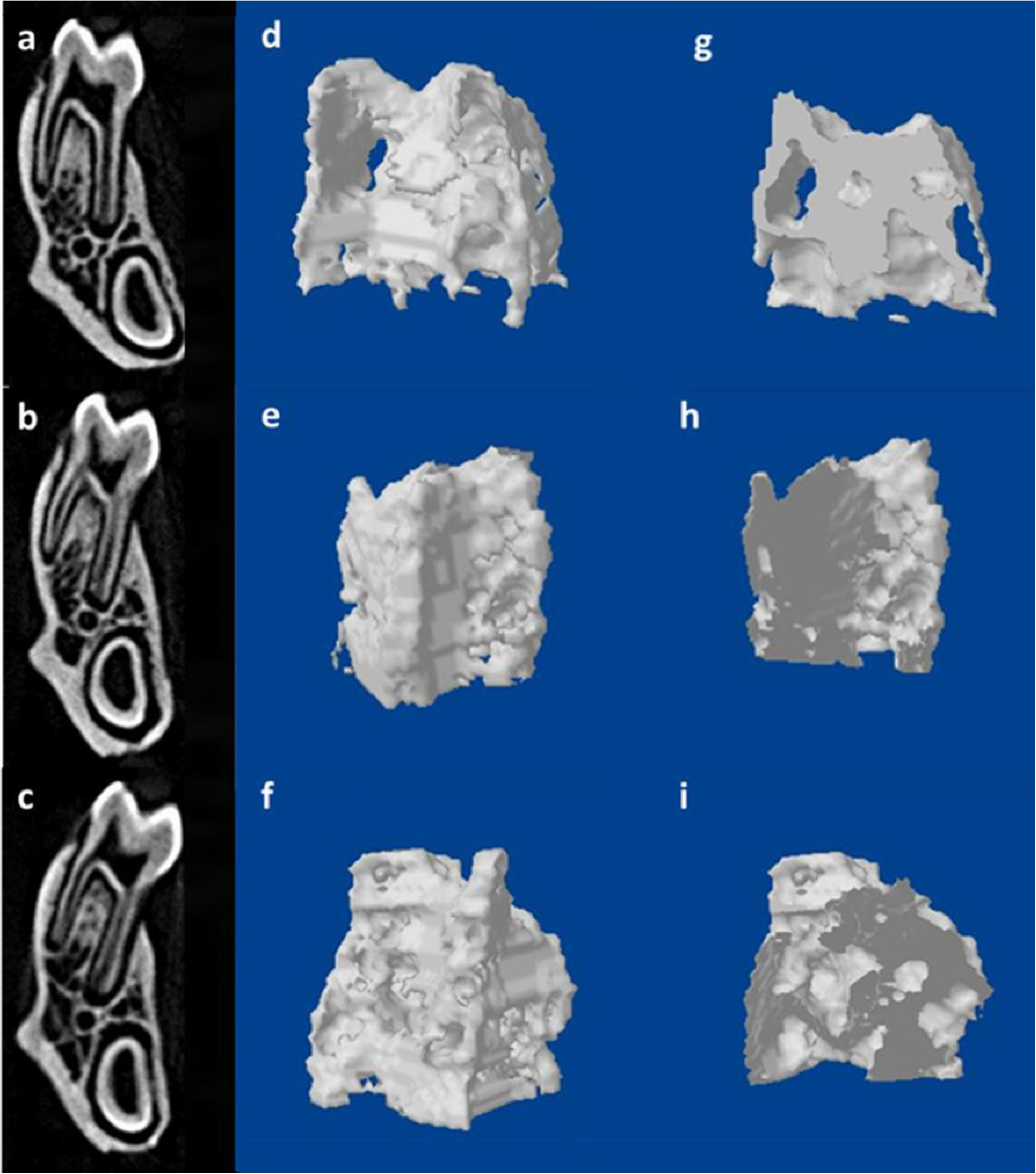
Organogram of the method used to obtain the three experimental groups: 1)CSO, Continuous Smoking Offsprings (in gray); 2)ISO, Interrupted Smoking Offsprings (in blue); and 3)NSO, Non-Smoking Offsprings or control (in white). Where is P, progenitors (in black) and SO, Smoking Offsprings (in red).

### Microtomography bone analysis

After the 21^st^, 42^nd^, 63^rd^ and 128^th^ days, the animals were sacrificed by anesthetic deepening (ketamine/xylazine) in accordance with the animal testing principles established by COBEA/CONCEA. After that, their mandibles were removed and stored in 10% buffered formaldehyde solution. Fifty-five samples were obtained (20 DTC, 20 DNT and 15 DTI) and scanned in a micro-computed tomography (SkyScan 1174, Kontich, Belgium), with 800 mA, 50kVp and time of exposure of 3800ms, for each rebuilt image and nominal isotropic resolution of 23μm established in the literature, according to Bouxsein et. al.[26]. It was used an aluminium filter (thickness 0,5mm). The scanning procedure lasted around 20 minutes, resulting in 270 images per sample. A global thresholding technique was used to binarize the mCT images in a gray scale where the minimum, between the apexes of the bone marrow and bone in the gray value histogram, was chosen with the limit value. The region of interest (ROI) of the alveolar bone was established manually in each image in the inter-root septum of the mandibular first molar (M1), on the right side, due to the concentration of tensions in this region, used in the histomorphometric analysis of the trabecular bone. [27]. The (ROI) was obtained according to the method used by Liu et al. (2015) that enabled the evaluation and visualization of the alveolar bone in an integral way [28].

At the beginning was identified, in bidimensional images, the coronal surface which passes through the midst of the intermediate buccal and lingual roots (Fig 2-a). After that, was chosen two horizontal surfaces, separeted one from each other, crossing the alveolar crest and the intermediate buccal apex (lines 1 and 2, Fig 2-parallel to the occlusion plane. The red portion shows the ROI in bidimensional image (Fig 2-a and 2-b). Finally, in a horizontal plan which showed the MI, was chosen the inter-root alveolar bone drawing an outline from the midst of a root canal to another, avoiding the roots and other structures in the bidimensional image. (Fig 2-b). After the reconstruction of the images in the software connected to the micro-computed tomography (NRecon SkyScan, Kontich, Belgium), the histomorphometry of the alveolar bone was obtained by the evaluation straight from the ROI 3D model, by the software of the system (CTAnalyser SkyScan, Kontich, Belgium). Were analyzed: the bone volume fraction (BV/TV), the trabecular thickness (Tb.Th), the trabeculae spaces (Tb.Sp), the trabecular number (Tb.N) and the Structure Model Index (SMI).

## Statistical analysis

Mean and standard deviation (SD) of the data generated by the software CTAnalyser were calculated by the software IBM-SPSS, version 17.0 (SPSS Inc., Chicago, IL, USA). It was also used the Shapiro-Wilk normality test and the Levene’s test of homogeneity of variances, with p<0,005. It was used Fisher’s variance analysis through the Gauss-Markov’s linear model and the multiple comparisons through the Tukey HSD’s criterion. Although the effect of the days compelled to repeat the measurements, the animals were sacrificed in each measurement in this experiment, being thus, different from a day to another. It was evaluated, to the different microtomographic analyzes, the error in the repetition of the measurements made by the same examiner. It was used the Dahlberg’s error formula in order to verify the intraobserver systematic error (apud HOUSTON, 1983)[29], with a significance level of 5%.

## Results

### Micro-computed tomography

The micro-computed tomography was used to investigate the microstructural alterations in the mandibular bone of all the offsprings in developing. Dahlberg’s formula about the intra-examiner systematic error (apud HOUSTON, 1983)[29] indicated a significance level of 4,1, validating the data collected. The trabecular parameters of the alveolar bone as the relation BV/TV, Tb.Th and SMI, showed significant differences in the CSO group, in relation to the values of the control group (Fig 4). Parameters such as Tb.Sp and Tb.N did not show significant differrences between the groups (Fig 4). In the DTC group, were observed significant reductions in BV/TV, p=0,0124, and inTb.Th, p<0,0001, associated to the significant increase of the SMI, p<0,0001, in relation to the control group (Table 1).

**Table 1.** The Microarchitectural Parametric Values, BV/TV, of the Mandibular Bone in the Inter-radicular Region (n=55)

| Group | Mean(%) and SD | Contrasts | P |
| --- | --- | --- | --- |
| NSO n20 | 69,35±16,69 | CSO / NOS | 0.0124 |
| CSO n20 | 56,81±13,33 | NSO / ISSO | 0.9751 |
| ISO n15 | 70,33±6,60 | CSO / ISSO | 0.0126 |

BV/TV: bone volume/ total volume; between CSO and NSO, and CSO and ISO $p<0,05$ ; between NSO and ISO, no significant values( $p>0,05$ ).

**Fig 3.**
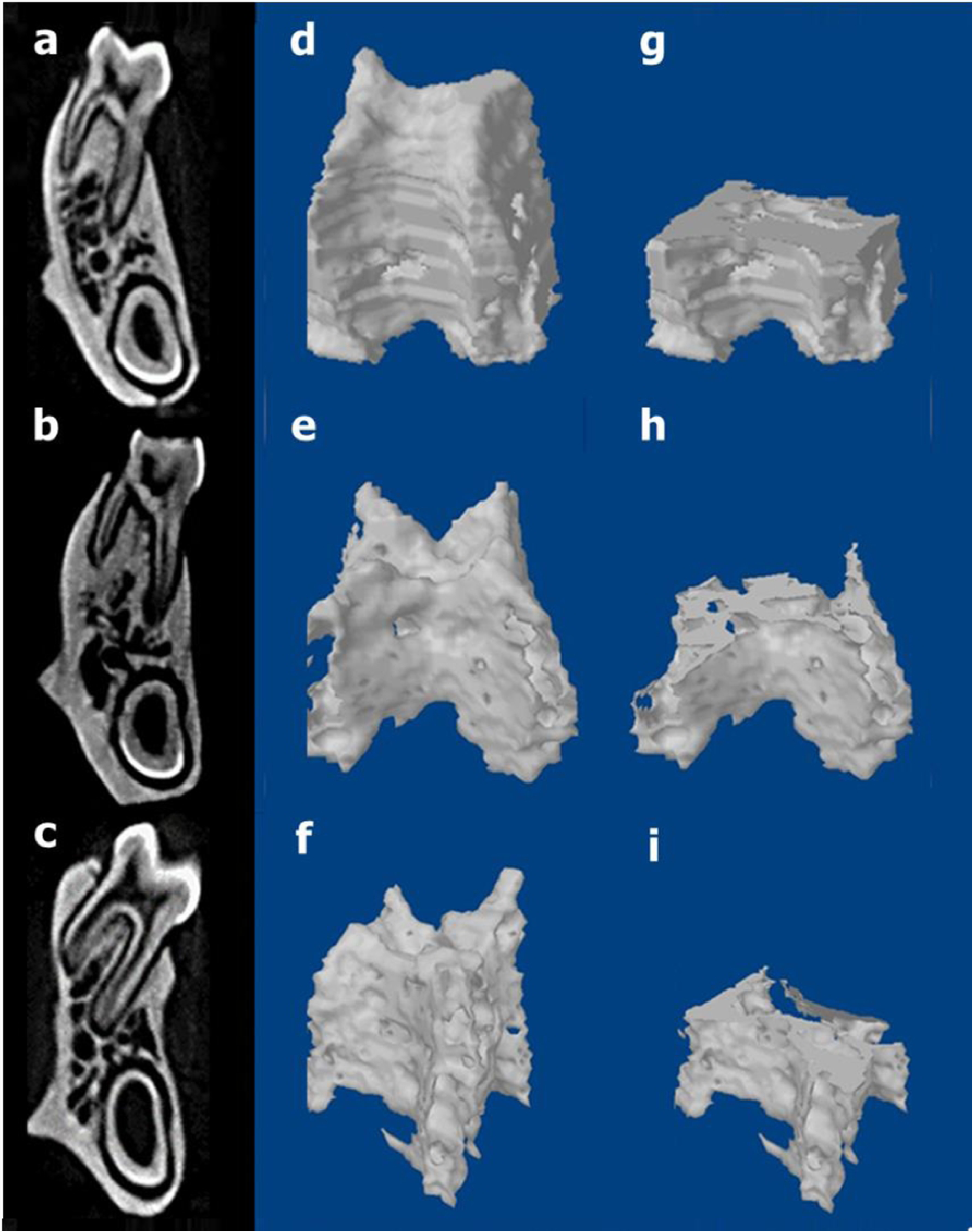
Micro-CT images of the rat mandibular alveolar bone and the mandibular first molar(M1); a) it shows the coronal surface passing through the midst of the intermediate buccal and lingual roots. The lines 1 and 2 pass by the alveolar crest and by buccal apex, respectively, parallel to the occlusion plane and delimiting the red area which represents the region of interest (ROI). B, buccal, L, lingual; **b)** horizontal surface parallel to the occlusion plane and the ROI between the midst of the four roots of the M1; M, mesial root; IB, intermediate buccal root; D, distal root; IL, intermediate lingual root.

**Fig 4.**
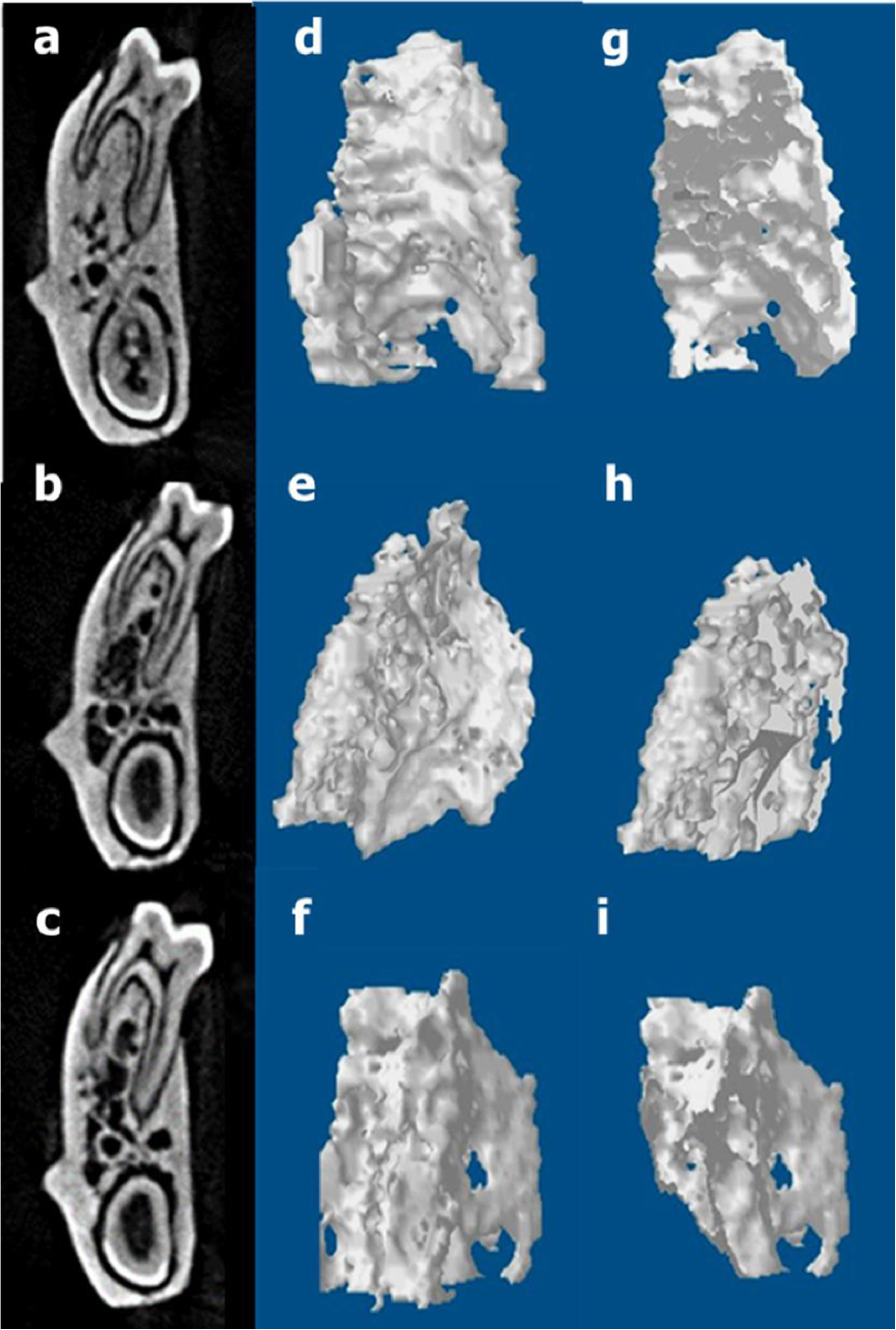
Micro-tomographic histomorphometric comparisons between the CSO (red), ISO (green) and NSO (blue) groups in the 21^st^, 42^nd^, 63^rd^ and 128^th^ days. A: BV/TV = bone volume fraction (bone volume/total volume); B: Tb.Th = trabecular thickness; C: SMI = Structure Model Index; D: Tb.Sp = trabecular spacing; E: Tb.N = trabecular number.

About the ISO group, there was no significant difference in the relation BV/TV, p=0,9751, when compared to the control group (Table 1). However, about Tb.Th, the differences among the three groups were significant, showing higher values in the control group, followed by the ISO group p=0,0020, and by the CSO group p=0,001, with the lower values (Table 2). The average values of the SMI for the ISO group, followed the values of the control group, p=0,1253, during the three periods studied (Table 3). The Tb.Sp did not show differences. It was not significant the effect of the exposure period to the cigarette smoke between the groups, p=0,6680 (Table 4). The Tb.N parameter, during all the exposure period to the cigarette smoke, there was no statistical difference between the groups p=0,3308 (Table 4). The values related to the trabecular bone alterations during all the experimental period, reveals about the CSO and NSO groups: difference of 18% in the BV/TV, 38,7% in the Tb.Th, and 51% in the SMI. Between CSO and ISO groups, 19,2% in the BV/TV, 24% in the Tb.Th, and 24,6% in the SMI. The difference between NSO and ISO was 1,4% in the BV/TV, 19,3% in the Tb.Th, and 14% in the SMI. The chart in the Fig 4 illustrates the tendencies of the mean for the three groups analysed, during the four periods. At the fig 5 – 8 its possible observing tomographic cross-sectional slices of the inter-radicular bone of M1 and representative images illustrating micro-CT image analysis in the inter-radicular region.

**Table 2.** The Microarchitectural Parametric Values, Tb.Th, of the Mandibular Bone in the Inter-radicular Region (n=55)

| Group | Mean(mm) and SD | Contrasts | P |
| --- | --- | --- | --- |
| NSO n20 | 0.31±0,05 | CSO / NOS | < 0.0001 |
| CSO n20 | 0.19±0,04 | NSO / ISSO | 0.0030 |
| ISO n20 | 0.25±0,05 | CSO / ISSO | 0.0020 |

Tb.Th: Trabecular thickness; statistical significant differences among the three groups, with $p < 0,005$ .

**Table 3.** The Microarchitectural Parametric Values, SMI, of the Mandibular Bone in the Inter-radicular Region (n=55)

| Group | Mean and SD | Contrasts | P |
| --- | --- | --- | --- |
| NSO n20 | 1.8±0,4 | CSO / NOS | < 0.0001 |
| CSO n20 | 2.7±0,2 | NSO / ISSO | 0.1253 |
| ISO n15 | 2.0±0,3 | CSO / ISSO | < 0.0001 |

SMI: Structure Model Index; between CSO and NSO, and CSO and ISO, $p < 0,05$ . Between NSO and ISO no significant values, with $p > 0,005$ .

**Table 4.** The Microarchitectural Parametric Values, Tb.Sp and Tb.N, of the Mandibular Bone in the Inter-radicular Region (n=55)

| Group | Tb.Sp Mean (mm) and SD | Tb.N Mean (mm) and SD |
| --- | --- | --- |
| NSO n20 | 0.2173±0,06 | 2,51±0,76 |
| CSO n20 | 0.2198±0,02 | 2,51±0,76 |
| ISO n15 | 0.1998±0,03 | 2,5707±0,51 |

Tb.Sp: trabecular spacing, TbN: trabecular number; no significant values to Tb.Sp, with $p = 0,6680$ and to Tb.N with $p = 0,3308$ .

**Fig 5.**
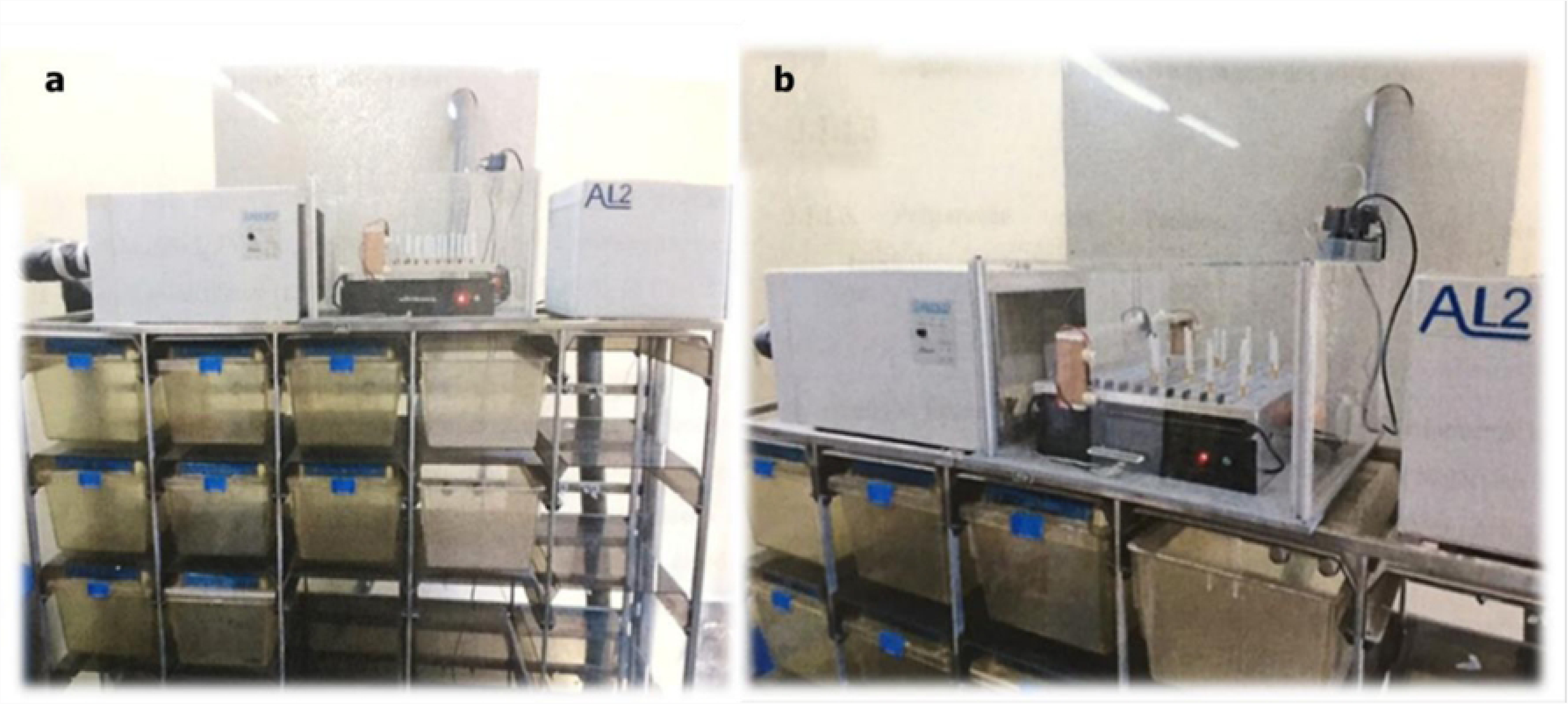
Tomographic cross-sectional slices of M1 furcation area(a,b) and representative images illustrating micro-CT image analysis in the inter-radicular region of the mandibular bone(c,d,e,f). a, c, e were obtained from a 21-day NSO rat, while b, d, f were obtained from a 21-day CSO rat. c, d showed the 3D reconstruction images, while e,f are the same images after section slice to observe the internal aspect. Images generated after the segmentation of osseous and non-osseous tissues demonstrated the ability to quantitatively analyze trabecular bone morphology

**Fig 6.**
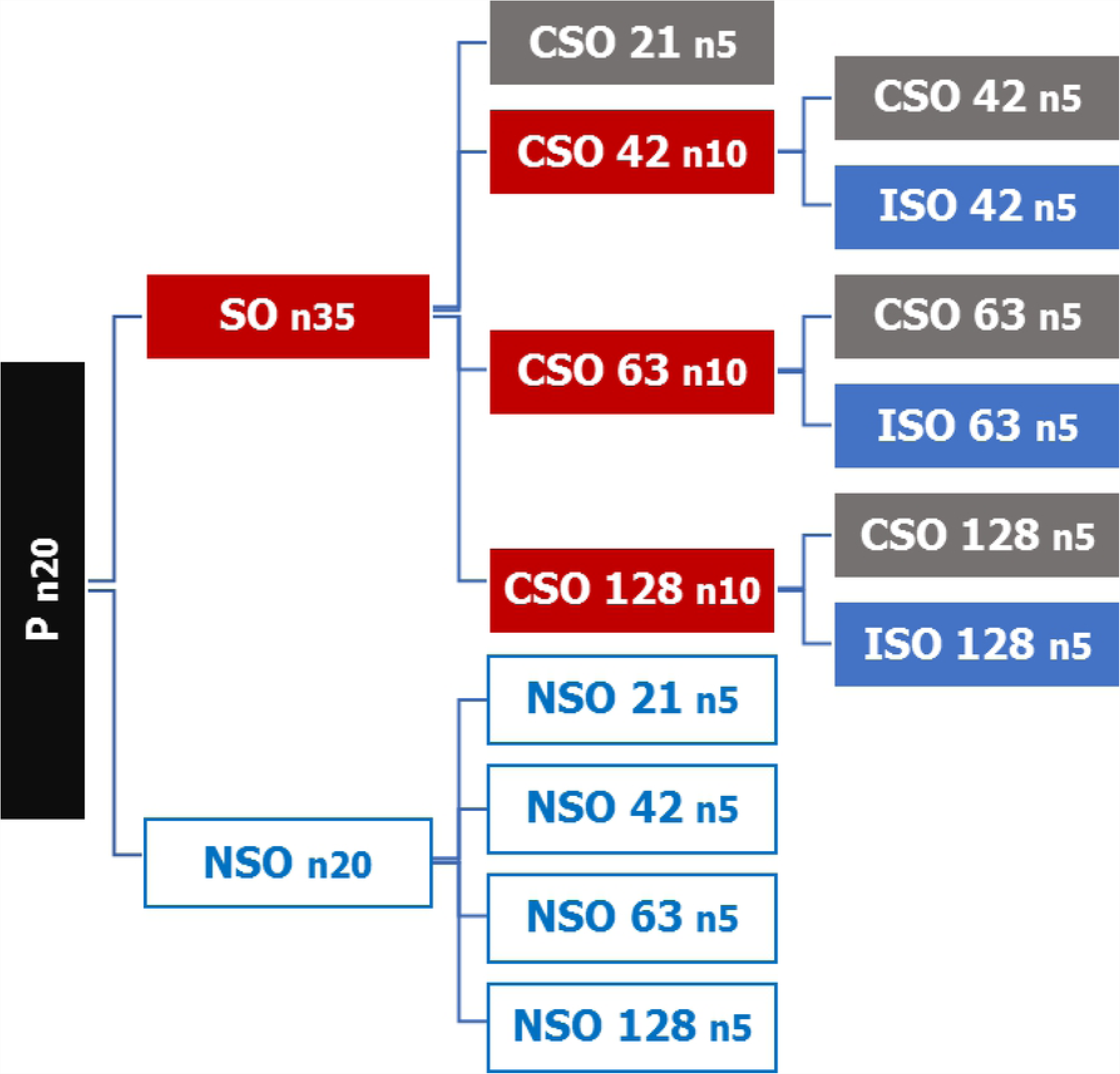
Tomographic cross-sectional slices of M1 furcation area(a,b,c) and representative images illustrating micro-CT image analysis in the inter-radicular region of the mandibular bone(d,e,f,g,h,i). a,d,g were obtained from a 42-day NSO rat, b,e,h were obtained from a 42-day ISO rat, and c,f,i were obtained from a 42-day CSO rat. d,e,f showed the 3D reconstruction images, while g,h,I are the same images after section slice to observe the internal aspect. Images generated after the segmentation of osseous and non-osseous tissues demonstrated the ability to quantitatively analyze trabecular bone morphology.

**Fig 7.**
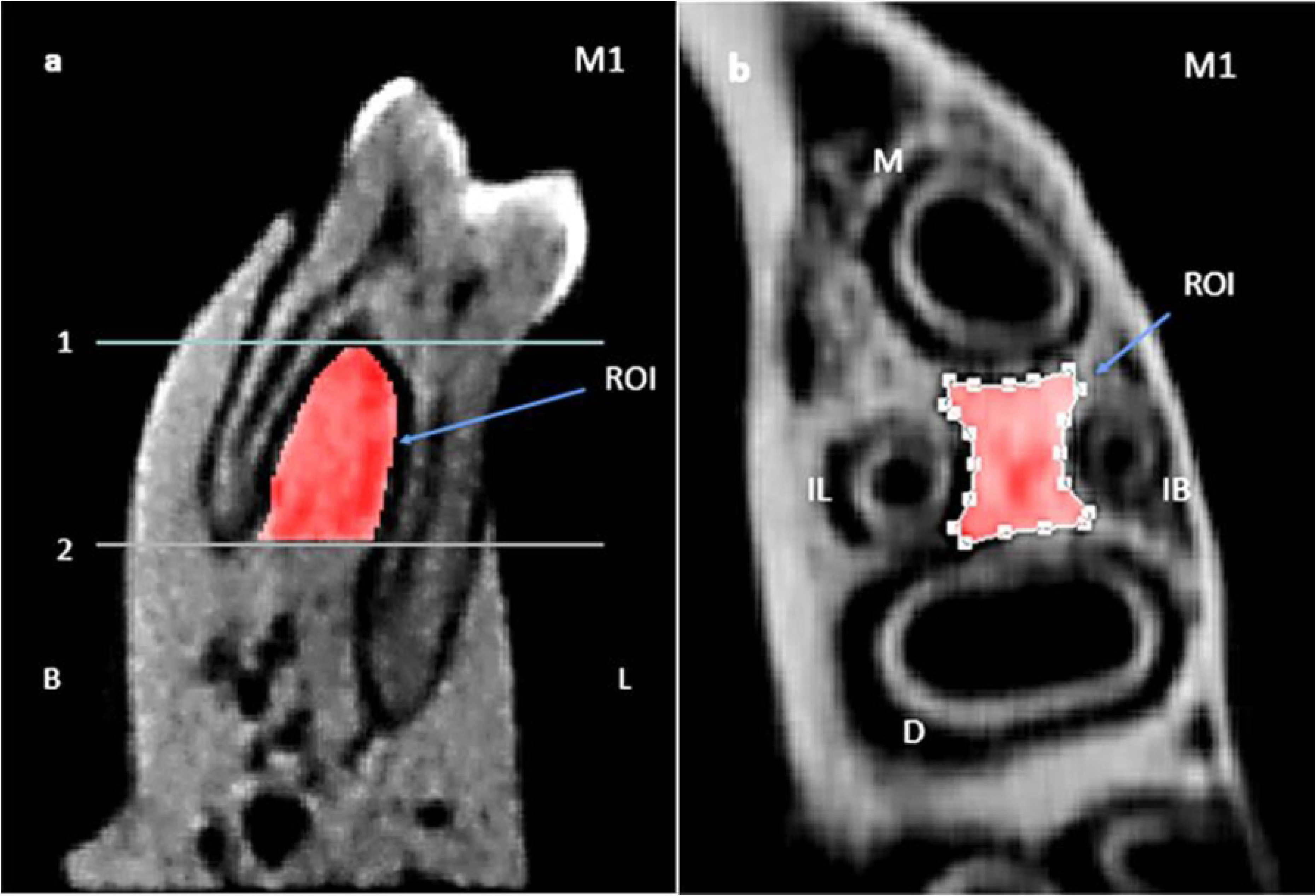
Tomographic cross-sectional slices of M1 furcation area(a,b,c) and representative images illustrating micro-CT image analysis in the inter-radicular region of the mandibular bone(d,e,f,g,h,i). a,d,g were obtained from a 63-day NSO rat, b,e,h were obtained from a 63-day ISO rat, and c,f,i were obtained from a 63-day CSO rat. d,e,f showed the 3D reconstruction images, while g,h,i are the same images after section slice to observe the internal aspect. Images generated after the segmentation of osseous and non-osseous tissues demonstrated the ability to quantitatively analyze trabecular bone morphology.

**Fig 8.**
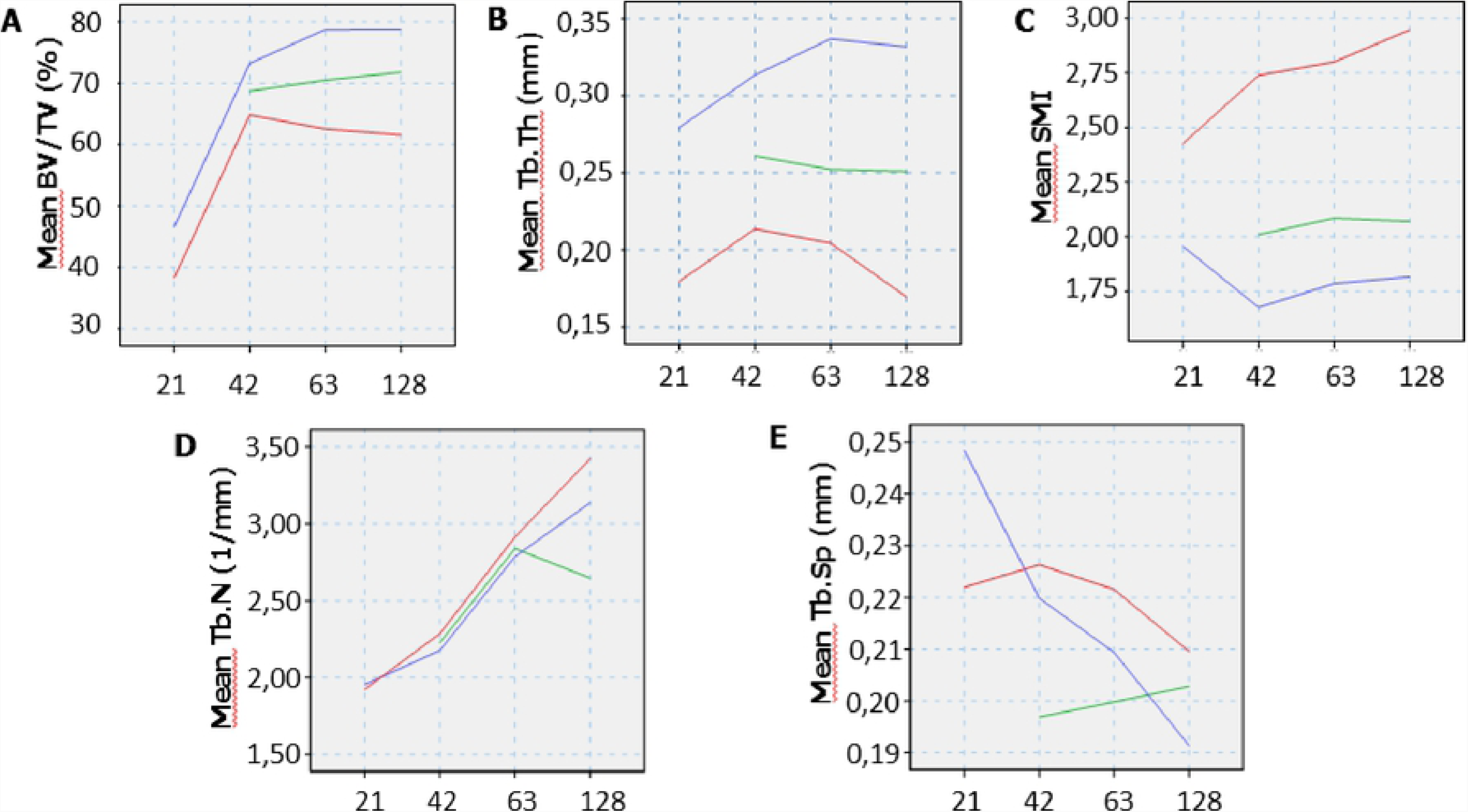
Tomographic cross-sectional slices of M1 furcation area(a,b,c) and representative images illustrating micro-CT image analysis in the inter-radicular region of the mandibular bone(d,e,f,g,h,i). a,d,g were obtained from a 128-day NSO rat, b,e,h were obtained from a 128-day ISO rat, and c,f,i were obtained from a 128-day CSO rat. d,e,f showed the 3D reconstruction images, while g,h,i are the same images after section slice to observe the internal aspect. Images generated after the segmentation of osseous and non-osseous tissues demonstrated the ability to quantitatively analyze trabecular bone morphology.

## Discussion

The present study simulated the environment of human beings when exposed to the cigarette smoke, similar to what happens in public smoking áreas and at home, where one or more of the family members are smokers. Our model was able to identify the secondhand smoke effects in the bone tissue of the animals evaluated in this study. The bone maturity of the Wistar rats happens from 119^th^ to 147 ^th^ days of life, that is equivalent, respectively, to the ages 18 and 24 in human beings [30]. In our study, was evaluated animals aged 21 days of life (that means 3 years old in humans), animals aged 42 days of life (means 7 years old in humans), animals aged 63 days of life (means 10 years old in humans), animals aged 128 days of life (means 21 years old in humans). Only the present study evaluated the characteristics of the bone tissue in development early periods, in offsprings of smoking matrices, considering also their effects after the cessation of the passive inhalation of the cigarette smoke.

The parameters BV/TV and Tb.Th, indicators of the trabecular bone density, show the development of the bone volume and of the trabecular thickness in the animals of the CSO group, from 21^st^ to 42^nd^ days of life. Showing exponential development, however, with lower values if compared with the control group. From 42^nd^ to 63^rd^ days of life, the rats undergo from puberty to sexual maturity [31], presented, tendency to loss bone density in the CSO group, that was intensified at the 128^th^ day. Dong et all. (2011)[32] showed general morphological inhibition in the development of the buccal tissues as a result of the maternal smoking, with lower values of thickness and volume in offsprings. Other studies showed the same significant reduction of the bone volume and trabecular thickness, indicating the suppression of the formation and increase of the bone resorption in rats exposed to the secondhand smoke. [19 -33 -34]

The ISO group showed recovery of the bone volume in relation to the CSO groups, however, the value of the trabecular thickness, between the animals of the CSO and control groups, was from 42^nd^ day on. That result suggests a weakened trabecular bone density. Reinforcing this finding, according to Hapidin et al. (2011), the cessation of the nicotine use during two months, does not allow the recovery of the adverse effects in the bone tissue, neither the return of the histomorphometric parameters at the control level [22]. Our results corroborate with the results of Ward et all (2001), who concluded that the cigarette smoke, in active smokers, has a harmful and dose-dependant effect causing the bone loss and increase of fracture risk and, that such effects are partially reversed due to the cessation of the inhalation [35]. The SMI, that was developed to indicate the relative prevalence of the elements in cylindrical and plate shapes in a 3D structure as the trabecular bone [36], is important in the evaluation of the osteoporotic degradation of the trabecular bone characterized by a transition from a plate architecture shape to a cylindrical architecture shape. The similar structure to the plates contributes to the bone elastic behavior, while the cylindrical shape results in a less rigid microstructure, and less able to withstand mechanical tensions coming from several directions [37]. In our study, there was a prevalence of cylindrical bone structural elements in the CSO group, that was increasing from 21^st^ to 128^th^ days, representing an increase of 51,1% in relation to the control. Ko et al. (2015), in their study, showed a similar result indicating that the secondhand smoke exposure increased in 46% the value of the SMI in relation to the control, confirming the osteopenic condition [1]. Sasaki M et al. (2018) showed in rats, that the secondhand smoke exposure, from gestation and birth to 28th day of life, increased the bone volume and induced to the osteoporosis, thanks to the spatial orientation of the tissue microstructure [38].

The analysis of the SMI in this study, indicated that the animals of the ISO group presented a surprising recovery in relation to the CSO groups. That shows the cessation of the inhalation causes the positive signaling to the osteoblastic cells, responsible for the bone formation. However, the trabecular thickness in intermediate levels, between the CSO and control groups, further suggests a fragile structure. Hapidin et al. (2011), showed that nicotine abstinente rats presented significant reduction in Tb.Th [21] in accordance with our data [22].

In our study, the offsprings of smoking matrices presented loss in the bone quality of the inter-radicular region (ROI) determined by the low values of BV/TV and Tb.Th. The values obtained were different from the values shown in previous studies [1-39], but the tendencies observed were consistent and the trabecular bone parameters, in the smoking group, were lower than in the control group. Our evaluation of the impact of secondhand smoke on bone development in rats from gestation to maturity has shown that detrimental effects on bone health increase with growth. We also showed that the offsprings removed from the exhibition, in half the time of the exposed animals, presented a favorable recovery, but under the level of control. This suggests a greater impairment in the bone recovery in the offsprings along the bone maturity and the effects of time exposure for which they were subjected to secondhand smoke exposured. Until the conclusion of this study, there are no reports about the evaluation of the toxic effects of the cigarette smoke, inhaled in a passive way, in other skeletal sites of the organisms in development. Thus, we concluded the need for an awareness of the Society about the toxic effects coming from the secondhand smoke exposure in the skeletal system of children and young people. Therefore, it is importante to avoid, in children and young people, whose parentes are smokers, the adverse effects in surgical intervations and healing processes. This care reduces the health risks in the face bones as well.

## Acknowledgments

The authors report no conflicts of interest related to this study. No external funding, apart from the support of the authors, was available for this study. Authors are greatful to Adriano Luis Martins and a Cristiano Manoel (State University of Campinas, Piracicaba, Department of Morphology) for their help in micro-CT analyses.

## Author contributions

### Conceptualization

CFA, CAFC, PHFC.

### Data curation

CFA.

### Formal analysis

CFA, AFI.

### Investigation

CFA, PHFC, CAFC, AFI.

### Methodology

CFA, CAFC, PHFC.

### Project administration

CFA, PHFC, CAFC.

### Supervision

PHFC, CAFC.

### Validation

PHFC, AFI, FHN.

### Visualization

CFA, PHFC, AFI, FHN.

### Writing ± original draft

CFA, PHFC.

### Writing ± review & editing

CFA, PHFC, CAFC.

## References

1. Ko CH, Chan RL, Siu WS, Shum WT, Leung PC, Zhang L, et al. Deteriorating effect on bone metabolism and microstructure by passive cigarette smoking through dual actions on osteoblast and osteoclast. Calcif Tissue Int. 2015 May;96(5):389–400

2. Schick S, Glantz S. Philip Morris toxicological experiments with fresh sidestream smoke: More toxic than mainstream smoke. TobControl 2005:14:396–404.

3. Website of Brazilian federal government (access in 2019 January 10) Available in: http://www.brasil.gov.br/saude/2011/05/males-do-fumo-passivo

4. Hirayama, T. (1981). Non-smoking wives of heavy smokers have a higher risk of lung cancer: a study from Japan. Brj Med J (Clin Res Ed), 282(6259), 183–185

5. Guo, S. & Dipietro, L. A. Factors affecting wound healing. J Dent Res. 2010;89(3), 219–29

6. Priemé H, Loft S, Klarlund M, Grønbaek K, Tønnesen P, Poulsen HE. Effect of smoking cessation on oxidative DNA modification estimated by 8-oxo-7,8-dihydro-2′-deoxyguanosine excretion. Carcinogenesis. 1998 Feb;19(2):347–51

7. Guo X, Oldham MJ, Kleinman MT, Phalen RF, Kassab GS. Effect of cigarette smoking on nitric oxide, structural, and mechanical properties of mouse arteries. Am J Physiol Heart Circ Physiol. 2006 Nov;291(5):H2354–61

8. Kanis JA, Johnell O, Oden A, Johansson H, De Laet C, Eisman JA, et al. Smoking and fracture risk: a meta-analysis. Osteoporos Int. 2005 Feb;16(2):155–62

9. Lee JJ, Patel R, Biermann JS, Dougherty PJ. The musculoskeletal effects of cigarette smoking. J Bone Joint Surg Am. 2013 May 1;95(9):850–9

10. Bouxsein ML. Bone quality: where do we go from here? Osteoporos Int. 2003 Sep;14 Suppl 5:S118–27.

11. Genant HK, Cooper C, Poor G, Reid I, Ehrlich G, Kanis J, et al. Interim report and recommendations of the World Health Organization Task-Force for Osteoporosis. Osteoporos Int. 1999;10(4):259–64.

12. Legrand E, Chappard D, Pascaretti C, Duquenne M, Krebs S, Rohmer V, Basle MF, Audran M. Trabecular bone microarchitecture, bone mineral density, and vertebral fractures in male osteoporosis. J Bone Miner Res. 2000 Jan;15(1):13–9.

13. Benowitz NL, Hall SM, Herning RI, Jacob P 3rd, Jones RT, Osman AL. Smokers of low-yield cigarettes do not consume less nicotine. N Engl J Med. 1983 Jul 21;309(3):139–42.

14. American Academy of Orthopaedic Surgeons (access in 2019 January 10) Available in: https://orthoinfo.aaos.org/en/staying-healthy/smoking-and-musculoskeletal-health/

15. Liu Z, Liu L, Kang C, Xie Q, Zhang B, Li Y. Effects of estrogen deficiency on microstructural change in rat alveolar bone proper and periodontal ligament. Mol Med Rep. 2015 Sep;12(3):3508–3514.

16. Wahl EA, Schenck TL, Machens HG, Egaña JT. Acute stimulation of mesenchymal stem cells with cigarette smoke extract affects their migration, differentiation, and paracrine potential. Sci Rep. 2016 Mar 15;6:22957.

17. Marinucci L, Bodo M, Balloni S, Locci P, Baroni T. Sub-toxic nicotine concentrations affect extracellular matrix and growth factor signaling gene expressions in human osteoblasts. J Cell Physiol. 2014 Dec;229(12):2038–48

18. Glowacki J, Schulten AJ, Perrott D, Kaban LB. Nicotine impairs distraction osteogenesis in the rat mandible. Int J Oral Maxillofac Surg. 2008 Feb;37(2):156–61

19. Ajiro Y, Tokuhashi Y, Matsuzaki H, Nakajima S, Ogawa T. Impact of passive smoking on the bones of rats. Orthopedics. 2010 Feb;33(2):90–5

20. Wirth AJ, Goldhahn J, Flaig C, Arbenz P, Müller R, van Lenthe GH. Implant stability is affected by local bone microstructural quality. Bone. 2011 Sep;49(3):473–8.

21. Inyang AF, Tchanque-Fossuo CN, Merati M, Radzolsky ER, Buchman SR. Microdensitometric and microarchitectural alterations in irradiated mandibular fracture repair. J Craniofac Surg. 2014 Nov;25(6):2022–6.

22. Hapidin H, Othman F, Soelaiman IN, Shuid AN, Mohamed N. Effects of nicotine administration and nicotine cessation on bone histomorphometry and bone biomarkers in Sprague-Dawley male rats. Calcif Tissue Int. 2011 Jan;88(1):41–7.

23. César-Neto JB, Benatti BB, Manzi FR, Sallum EA, Sallum AW, Nociti FH. The influence of cigarette smoke inhalation on bone density. A radiographic study in rats. Braz Oral Res. 2005 Jan-Mar;19(1):47–51

24. Santiago HA, Zamarioli A, Sousa Neto MD, Volpon JB. Exposure to Secondhand Smoke Impairs Fracture Healing in Rats. Clin Orthop Relat Res. 2017 Mar;475(3):894–902

25. Müller R, Van Campenhout H, Van Damme B, Van Der Perre G, Dequeker J,Hildebrand T, et al. Morphometric analysis of human bone biopsies: a quantitative structural comparison of histological sections and micro-computed tomography. Bone. 1998 Jul;23(1):59–66.

26. Bouxsein ML, Boyd SK, Christiansen BA, Guldberg RE, Jepsen KJ, Müller R. Guidelines for assessment of bone microstructure in rodents using micro-computed tomography. J Bone Miner Res. 2010 Jul;25(7):1468–86. doi:10.1002/jbmr.141. Review. PubMed PMID:20533309.

27. Shimizu Y, Ishida T, Hosomichi J, Kaneko S, Hatano K, Ono T. Soft diet causes greater alveolar osteopenia in the mandible than in the maxilla. Arch Oral Biol. 2013 Aug;58(8):907–11.

28. Liu Z, Yan C, Kang C, Zhang B, Li Y. Distributional variations in trabecular architecture of the mandibular bone: an in vivo micro-CT analysis in rats. PLoS One. 2015 Jan 27;10(1):e0116194.

29. Houston, W. J. B. The analysis of errors in orthodontic measurements. Am. J. Orthod. 1983 May; St. Louis. 83(5):382–90.

30. Fukuda S, Matsuoka O. Maturation process of secondary ossification centers in the rat and assessment of bone age. Jikken Dobutsu. 1979 Jan;28(1):1–9.

31. Andrade A, Pinto S. C, de Oliveira R. S. Laboratory animals: breeding and experimentation. Rio de Janeiro, Rio de Janeiro. FIOCRUZ; 2002.

32. Dong Q, Wu H, Dong G, Lou B, Yang L, Zhang L. The morphology and mineralization of dental hard tissue in the offspring of passive smoking rats. Arch Oral Biol. 2011 Oct;56(10):1005–13.

33. Wang D, Nasto LA, Roughley P, Leme AS, Houghton AM, Usas A, et al. Spine degeneration in a murine model of chronic human tobacco smokers. Osteoarthritis Cartilage. 2012 Aug;20(8):896–905

34. Gao SG, Cheng L, Li KH, Liu WH, Xu M, Jiang W, et al. Effect of epimedium pubescen flavonoid on bone mineral status and bone turnover in male rats chronically exposed to cigarette smoke. BMC Musculoskelet Disord. 2012 Jun 19;13:105

35. Ward KD, Klesges RC. A meta-analysis of the effects of cigarette smoking on bone mineral density. Calcif Tissue Int. 2001 May; 681:259–70

36. Hildebrand T, Rüegsegger P. Quantification of Bone Microarchitecture with the Structure Model Index. Comput Methods Biomech Biomed Engin. 1997;1(1):15–23.

37. Liu XS, Sajda P, Saha PK, Wehrli FW, Guo XE. Quantification of the roles of trabecular microarchitecture and trabecular type in determining the elastic modulus of human trabecular bone. J Bone Miner Res. 2006 Oct;21(10):1608–17

38. Sasaki M, Chubachi S, Kameyama N, Sato M, Haraguchi M, Miyazaki M,et al. Effects of long-term cigarette smoke exposure on bone metabolism, structure, and quality in a mouse model of emphysema. PLoS One. 2018 Jan 30;13(1):e0191611.

39. Bakathir MA, Linjawi AI, Omar SS, Aboqura AB, Hassan AH. Effects of nicotineon bone during orthodontic tooth movement in male rats. Histological and immunohistochemical study. Saudi Med J. 2016 Oct;37(10):1127–35.

